# Immunosuppressive role of PGE2 during human tuberculosis

**DOI:** 10.1101/2020.08.04.236257

**Authors:** Joaquín Miguel Pellegrini, Nancy Liliana Tateosian, María Paula Morelli, Agustín Rollandelli, Nicolás Oscar Amiano, Domingo Palmero, Alberto Levi, Lorena Ciallella, María Isabel Colombo, Verónica Edith García

## Abstract

Prostaglandin E2 (PGE2), an active lipid compound derived from arachidonic acid, regulates different stages of the immune response of the host during several pathologies such as chronic infections or cancer. Manipulation of PGE2 levels was proposed as an approach for countering the Type I IFN signature of tuberculosis (TB), but very limited information exists about this pathway in patients with active TB. Here, we demonstrated that PGE2 exerts a potent immunosuppressive action during the immune response of the human host against *M. tuberculosis*. Thus, we showed that PGE2 inhibited both lymphoproliferation and cytokine production of proinflammatory cytokines, together with a significant reduction of the surface expression of several immunological receptors in human cells. However, PGE2 promoted the autophagic flux of antigen-stimulated monocytes, even in the presence of IFNα. In this way, the attenuation of inflammation and immunopathology caused by an excessive immune response emerges as an attractive therapeutic target. Together, our findings contribute to the knowledge of *Mtb*-resistance mediated by PGE2 and highlight the potential of this lipid mediator as a tool to improve anti-TB treatment.

## Introduction

It is estimated that *Mycobacterium tuberculosis* (*Mtb*), the etiologic agent of tuberculosis (TB), has killed nearly 1000 million people in the last two centuries^1^. Furthermore, nowadays, TB remains a major global health problem ranking among the top ten causes of death worldwide. Despite the availability of an affordable and effective treatment, it is the leading cause of death from a single infectious agent, accounting for about a quarter of preventable mortality in developing countries^2^. Thus, the main goals of developing new strategies to fight *Mtb* include to reduce the treatment length, to address the drug resistance’s problems, to provide safer treatments and to eradicate the drug interactions observed in patients with HIV/TB infection^3^.

Although great progress has been made in characterizing the immune response acquired in TB patients, it remains to be elucidated what are the most appropriate defense mechanisms required to fight *Mtb*. The immune response of the host against Mtb is highly complex and the outcome of the infection depends on immunological mediators and their particular temporal dynamics in the microenvironment. In this regard, in addition to the crucial role of cytokines and chemokines, accumulated evidence in recent years has indicated that different lipid mediators, mainly eicosanoids, are critical in the resolution of the mycobacterial infection.

Eicosanoids are a family of bioactive lipid mediators derived from arachidonic acid (AA), which is released from membrane phospholipids by phospholipases. There are three main pathways involved in the production of eicosanoids, which typically compete with each other for AA: (i) the cyclooxygenase pathway (COX-1 and COX-2) that produces prostaglandins and thromboxanes, (ii) the pathway of lipoxygenases (5-LOX, 12-LOX, and 15-LOX), that catalyzes the production of leukotrienes and lipoxins, and (iii) the cytochrome p40 pathway, which generates hydroxyeicosatetraenoic and epoxieicosatrienoic acids^4^.

The ensuing findings that prostaglandin E (PGE) synthase-deficient mice^5^ and mice lacking the prostaglandin receptor EP2 have increased susceptibility to *Mtb* infection^6^ provide strong evidence that PGE2 and the apoptotic death of macrophages are critical to regulate *Mtb* growth *in vivo*^5^, although the exact mechanisms of this protection have not been elucidated. In this regard, Chen *et al*. reported that avirulent *Mtb* strain H37Ra induces PGE2 production, which protects against cell necrosis by preventing the internal mitochondrial membrane damage^5^ and promoting a rapid plasma membrane repair^7^. Conversely, PGE2 is immunosuppressive for T cell-mediated immunity at high concentrations^8^ and contributes to the expansion of regulatory T cells^9^ during infection with *Mtb*, although its exact role during adaptive immunity in human TB is uncertain. In an infection model using IL-1 signaling deficient mice or animals that produced excessive type I IFN, administration of both zileuton and PGE2 showed host-beneficial effects^10^. Inhibition of the enzyme 5-lipooxygenase (5-LO) is proposed to shunt eicosanoid production toward increased PGE2 production, which in the context of high-inflammatory environments might suppress detrimental type I IFN production, partially rescue mortality, and reduce bacterial burden^10^. Thus, manipulation of PGE2 and/or 5-LO could serve to counteract these detrimental responses in patients with severe TB. However, the precise mechanism of PGE2 and their impact on the immune response of the host during TB remain to be studied. Importantly, the ultimate goal of those studies will open new avenues to develop host-targeted therapies. Hence, it is crucial the functional characterization of the soluble mediators that participate in the immune response against *Mtb*, in order to achieve an improvement in the design of vaccines and joint immunotherapies that considerably improve control over TB.

Therefore, in this work we studied the role of PGE2 in the immune response of the host during human TB infection. Our main goal was to determine the cellular and physiological bases of PGE2 function to evaluate it as part of a potential host directed therapy.

## Results

First, plasma levels of PGE2 were determined in peripheral blood from healthy donors (HD) and TB patients by radioimmunoassay. As shown in Figure 1, and as previously reported by others^10^, we found significantly higher concentrations of PGE2 in the plasma of patients with TB as compared to HD, denoting the inflammatory state of the infected individuals. It is important to mention that a small percentage of the recruited TB patients (15%) were under treatment with ibuprofen, a common medication given to these patients to reduce fever and other the symptoms of the disease. In these patients, we detected a lower plasmatic concentration of PGE2 in comparison with non-treated TB patients (data not shown), as expected due to the inhibitory function of ibuprofen on COX-2 enzyme activity.

**Figure 1.**
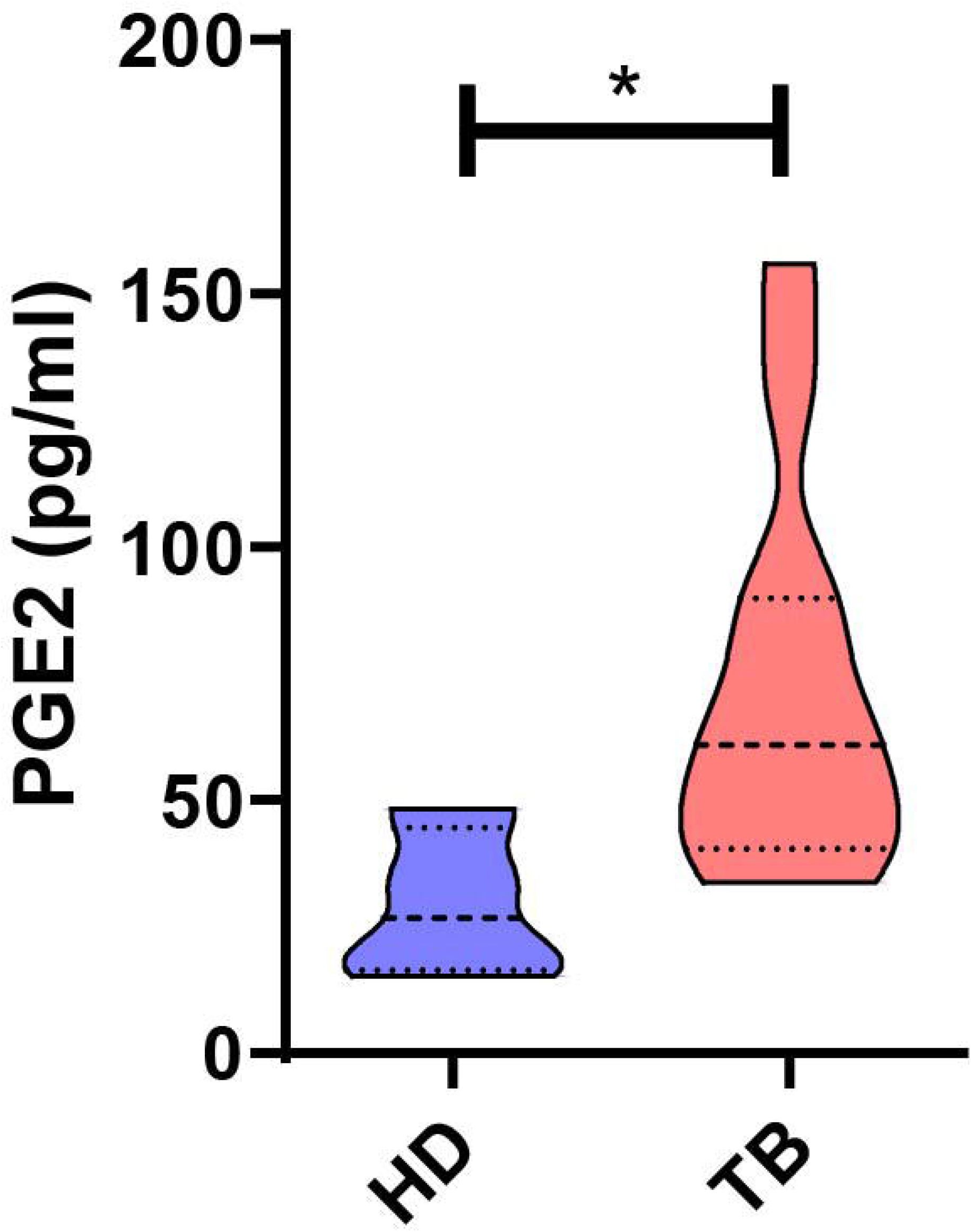
PGE2 Plasma levels from Healthy Donors and TB patients. Heparinized peripheral blood from healthy donors (HD, n = 5) and TB patients (n = 10) was centrifuged for 15 minutes at 1000g and the levels of PGE2 in plasma were analyzed by radioimmunoassay. Bars show the mean values of PGE2 plasma concentration (pg/ml) ± SEM. P values were calculated using the Mann Whitney non-parametric test for unpaired samples. * p <0.05

In order to further study the role of PGE2 on the adaptive immune response of the human host against *Mtb* infection, we next investigated the effect of this eicosanoid on lymphocyte proliferation in HD and TB patients. Thus, PBMC from both groups of individuals were stimulated for 5 days with a *Mtb* lysate (*Mtb*-Ag) in the presence or absence of PGE2 at different concentrations. Those concentrations were similar or higher to PGE2 levels found at the infection site during specific treatments. Finally, cell proliferation was analyzed by tritiated thymidine incorporation over the last 16 hours of culture. As shown in Figure 2A, even at low concentrations of PGE2, a significant inhibition of *Mtb*-Ag induced lymphoproliferation was observed compared to cells only stimulated with *Mtb*-Ag. Furthermore, this suppressive effect of PGE2 on cell proliferation was verified in cells from both HD and TB patients (Figure 2B).

**Figure 2.**
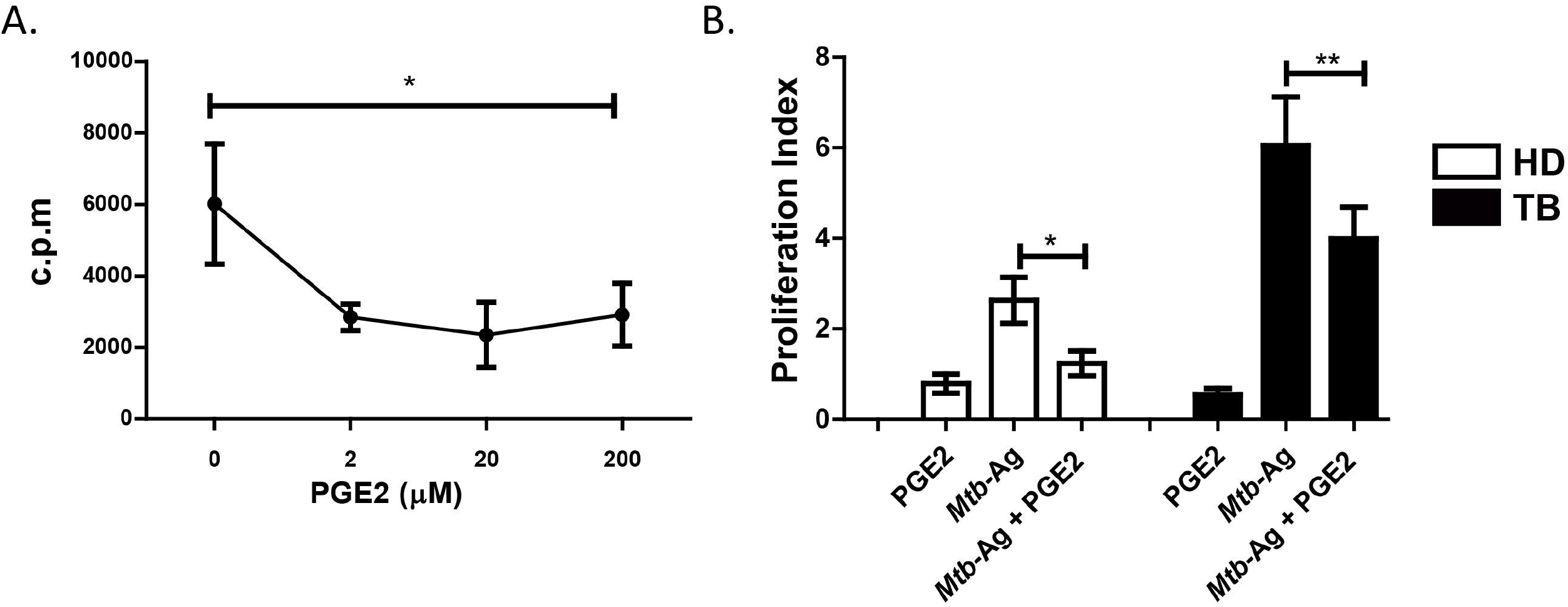
PGE2 inhibits lymphocyte proliferation. **(A)** Dose-response curve of PGE2 (0-200 μM) treatment and effect on proliferation of Peripheral blood mononuclear cells (PBMC) from HD stimulated with *Mtb*-Ag (10 μg/ml). **(B)** PBMC from healthy donors (HD) and TB patients were stimulated with *Mtb*-Ag (10 μg/ml) in the presence or absence of PGE2 (2 μM) for 5 days. Afterwards, cellular proliferation was analyzed by incorporation of Titrated thymidine during the last 16h of culture. Bars show the values of proliferation index (c.p.m. in each condition/c.p.m. media) ± SEM. Statistical differences were calculated using one-way ANOVA and post hoc Dunnett multiple comparison test. *p<0.05, ** p<0,01.

Subsequently, we decided to evaluate the expression of surface receptors involved in T lymphocyte activation in the presence of PGE2. To do this, PBMC from HD and TB patients were stimulated with *Mtb*-Ag in the presence or absence of PGE2 for 5 days, and then the expression of several immune receptors was evaluated by flow cytometry.

Interestingly, we observed that PGE2 treatment significantly reduced SLAMF1 and CD31 expression on CD3^+^ T lymphocytes (Figure 3A, B). Moreover, a significant decreased in the expression of the costimulatory molecule CD80 and MHC class I was also confirmed on CD14^+^ monocytes treated with this eicosanoid (Figure 3C, D). Besides, we also detected a decreasing trend in MHC class II expression on cells stimulated with *Mtb* antigen in the presence of PGE2 (Figure 3E). Taken together, these results suggest that PGE2 might participate in the regulation of T cell activation in response to mycobacterial antigens.

**Figure 3.**
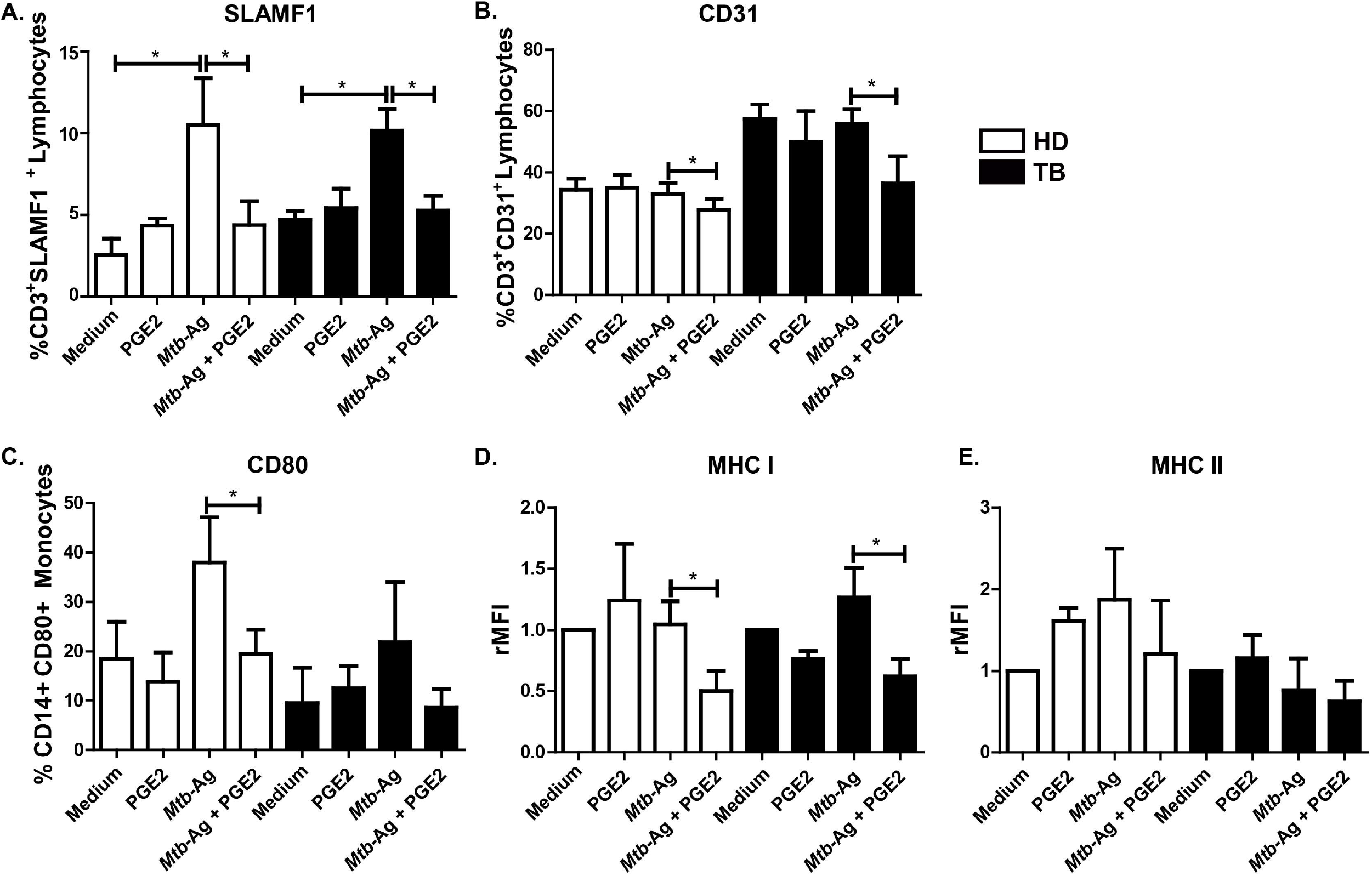
PGE2 regulates surface receptor expression on PBMC from HD and TB patients. Peripheral blood mononuclear cells (PBMC) from HD and TB patients were stimulated with *Mtb*-Ag (10 μg/ml) in the presence or absence of PGE2 (2 μM) during 5 days. Afterwards, the expression of **(A)** SLAMF1, **(B)** CD31 on CD3^+^ T cells and **(C)** CD80, **(D)** MHC-I and **(E)** MHC-II on CD14^+^ monocytes were determined by flow cytometry. Each bar represents **(A, B, C)** the mean percentage of CD14^+^ or CD3^+^ Receptor^+^ cells ± SEM or **(D, E)** relative Mean Fluorescence Intensity (rMFI) ± SEM. Statistical differences were calculated using one-way ANOVA and post hoc Dunnett multiple comparison test. *p<0.05.

Additionally, we analyzed the effect of PGE2 treatment on the production of key cytokines that participate in the immune response against *Mtb*. Therefore, we investigated the levels of IFNγ, IL-17A, TNFα and IL-1β in PBMC culture supernatants stimulated with *Mtb*-Ag in the presence or absence of PGE2. In line with the results described above, PGE2 induced a significant decrease in IFNγ and TNFα production in response to *Mtb*-Ag (Figure 4A, B). Moreover, PGE2 also markedly decreased IL-1 β and IL-17A, although no significant differences were detected (Figure 4C and data not shown).

**Figure 4.**
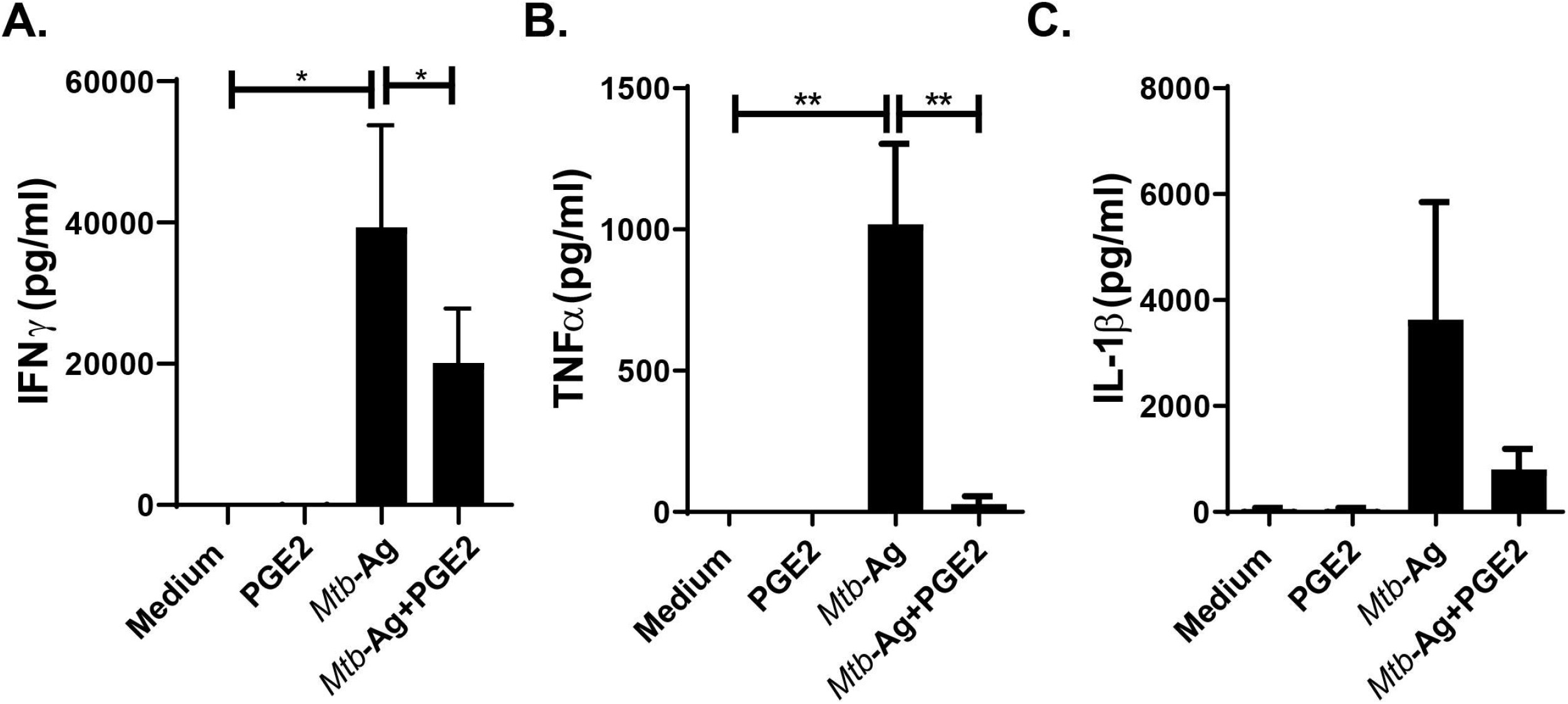
PGE2 modulates cytokine production. PBMC from TB patients were stimulated with *Mtb*-Ag (10 μg/ml) in the presence or absence of PGE2 (2 μM) for **(A)** 5 days or **(B, C)** 24 h. Afterwards, the production of **(A)** IFNγ, **(B)** TNFα and **(C)** IL-1β were determined by ELISA. Each bar represents the mean of cytokine production (pg/ml) ± SEM. Statistical differences were calculated using one-way ANOVA and post hoc Dunnett multiple comparison test. *p<0.05, ** p<0.01.

Taken together, the results presented so far suggested a suppressive/regulatory role for PGE2 on the innate and adaptive immune responses against *Mtb*. However, in contrast to our findings, different studies have shown a beneficial role for PGE2 during infection with *Mtb*^5^. Furthermore, PGE2 is proposed as a host-directed treatment to counteract the predominant responses of type I interferons displayed by severe TB patients^10^. The PGE2-dependent cellular mechanisms that operate on infected macrophages have not been fully elucidated. Importantly, previous studies observed that PGE2 protects against cell necrosis by preventing damage to the internal mitochondrial membrane^5^ and promoting rapid repair of the plasma membrane^7^.

In this context, autophagy arises as a potential mechanism that may explain the positive role of this eicosanoid during the early stages of *Mtb* infection. In this regard, it has been shown that cyclic mechanical stretching induces autophagic cell death in tenofibroblasts through the activation of PGE2production^11^ and that nicotine induces cell stress and autophagy mediated by COX-2 activation and PGE2 production in human colon cancer cells^12^. Furthermore, it was recently described that the expression of COX-2 contributes to increase autophagy and therefore to eliminate *Mtb* in bone marrow murine derived macrophages^13^. Nevertheless, this modulation has not been described yet in primary human cells. Therefore, we next analyzed autophagy modulation by PGE2 in *Mtb*-Ag stimulated-PBMCs from HD and TB patients.

First, we performed a kinetic study in which the autophagy levels in stimulated-cells from HD and TB patients were evaluated by flow cytometry as described before^14^. As shown in Figure 5A and B, a significant increase in the percentage of CD14^+^LC3A, B-II^+^ monocytes and in the relative mean fluorescence intensity (rMFI) was detected in cells treated with PGE2 for 16 hours. However, the augmented levels of autophagy decreased to those of untreated cells at 24 hours (Figure 5A, B). The augmented LC3A, B-II levels were corroborated both in monocytes from HD and TB patients (Figure 5C), while a not significant similar trend was observed in lymphocytes of both populations of individuals (Figure 5D). Likewise, confocal microscopy analysis yielded similar results, denoted by a greater number of LC3 puncta per cell in *Mtb*-Ag stimulated-monocytes treated with PGE2 as compared to untreated cells (Figure 6 E, F).

**Figure 5.**
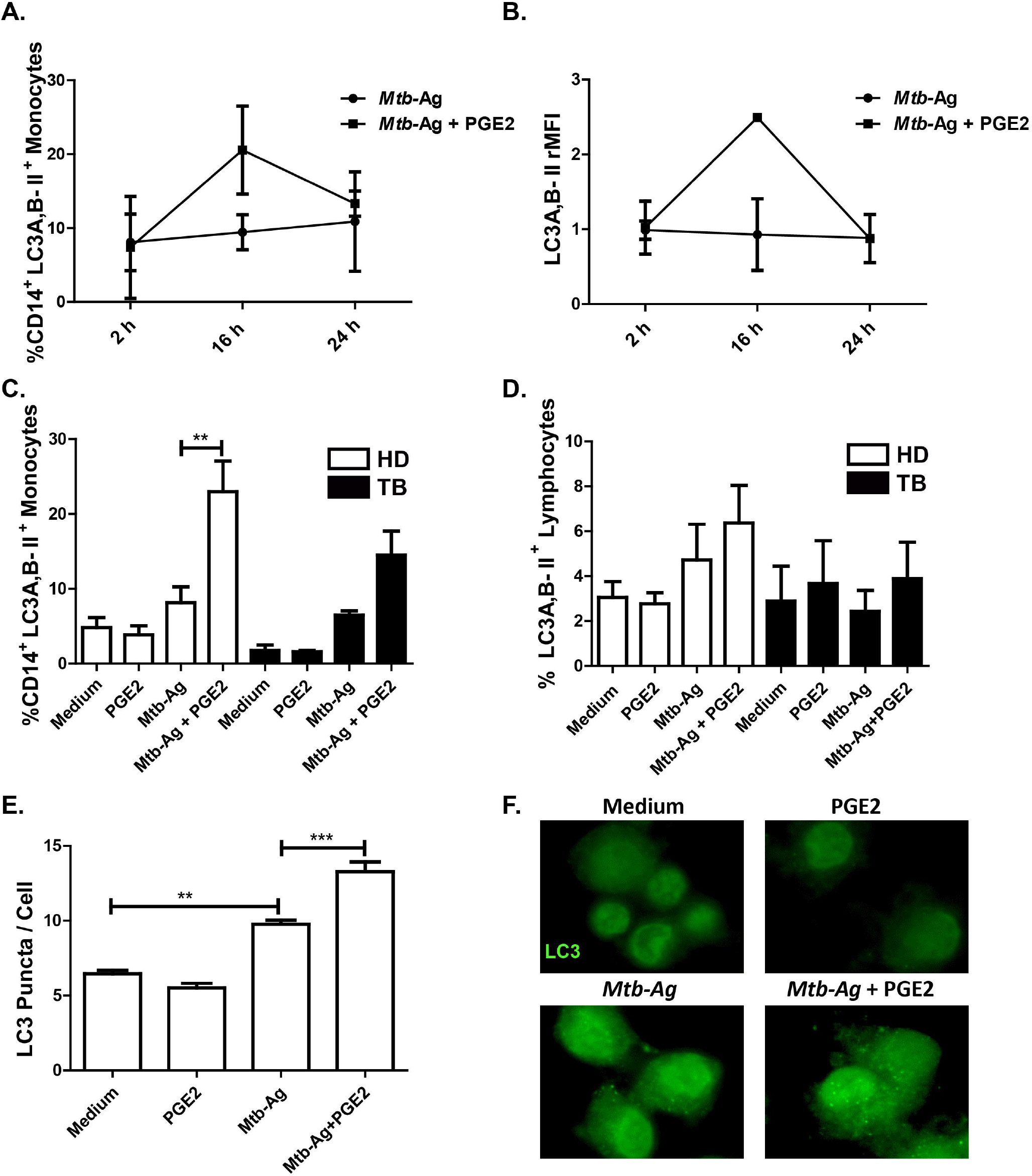
PGE2 induces autophagy in *Mtb*-Ag-stimulated cells from healthy donors and TB patients. Adherent cells from healthy donors (HD) and TB patients were stimulated with *Mtb*-Ag (10 μg/ml) in the presence or absence of PGE2 (2 μM) at the indicated time points. The levels of autophagy were then evaluated by intracellular flow cytometry through an indirect protocol using anti-LC3A, B-II mAb in CD14^+^ cells. **(A)** Percentage of CD14^+^LC3-IIB^+^ cells ± SEM; **(B)** Mean fluorescence intensity (MFI) of CD14^+^LC3A, B-II^+^ cells ± SEM. **(C,D)** Bars represent the percentage of **(C)** monocytes CD14^+^ LC3A, B-II^+^ and **(D)** lymphocytes LC3A, B-II^+^ after 16h of stimulation. **(E)** PBMC from HD were incubated at 2×10^6^ cells/ml during 16h to allow adherence of monocytes. Afterwards, the cells were stimulated with *Mtb*-Ag (10 μg/ml) in the presence or absence of PGE2 (2 μM) during 16h. Autophagy levels were evaluated by immunofluorescence against LC3 on monocytes. Bars represent the mean values of LC3 puncta per cell ± SEM. **(F)** A representative experiment is shown. Bars represent the mean values of the percentage of LC3A, B-II^+^ cells ± SEM. Statistical differences were calculated using one-way ANOVA and post hoc Dunnett multiple comparison test. *p<0.05, ** p<0.01, *** p<0.001.

**Figure 6.**
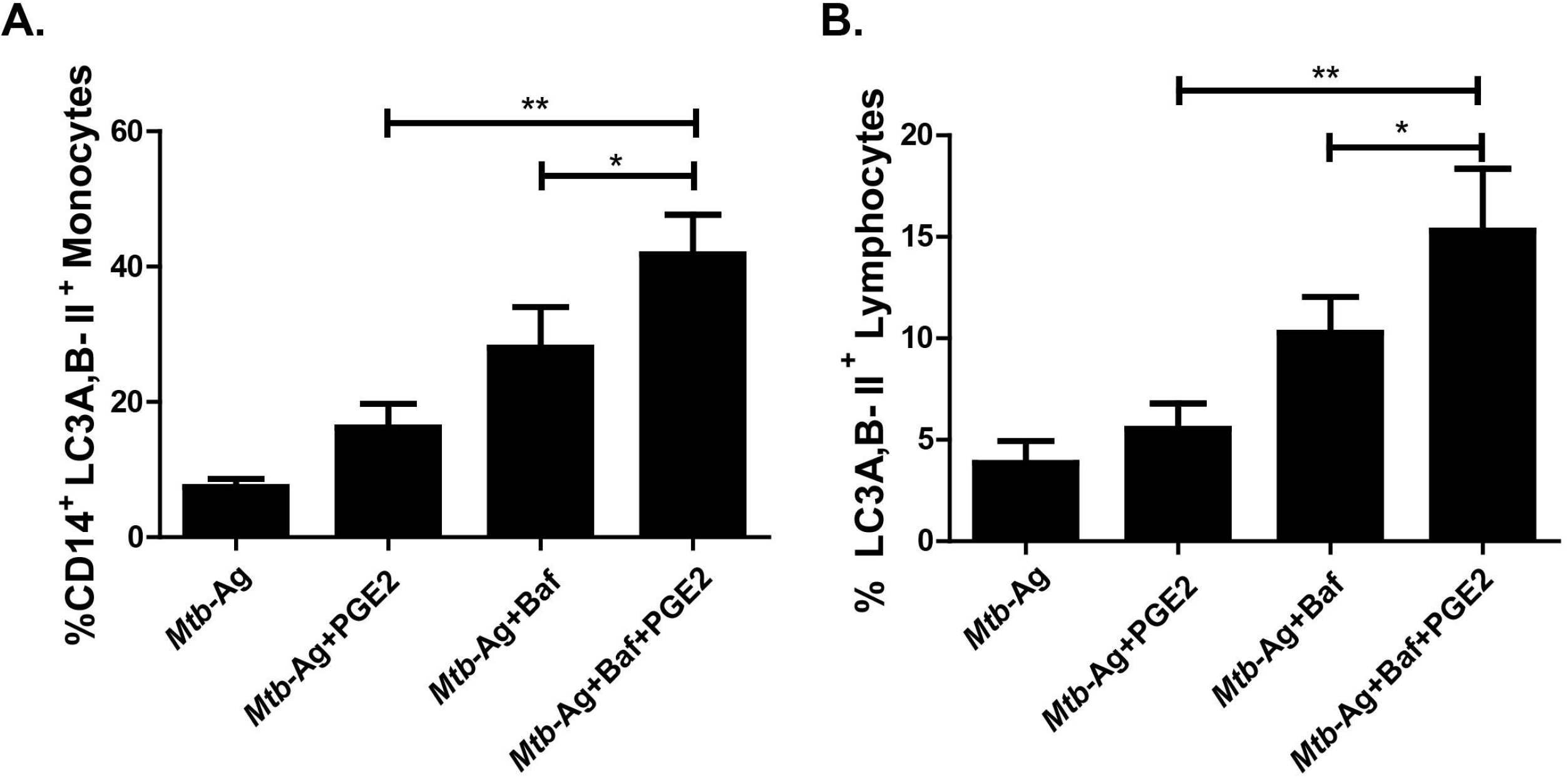
Autophagy flow in cells treated with PGE2. PBMC from TB patients were incubated at 2×10^6^ cells/ml during 16h to allow adherence of monocytes. Afterwards, the cells were stimulated with *Mtb*-Ag (10 μg/ml) in the presence or absence of PGE2 (2 μM) during 16h. To evaluate autophagy flow, Bafilomycin A1 (Baf A1, 1 μg / ml) was added in the last 2h of culture before determining the autophagy levels by flow cytometry by means of indirect intracytoplasmic staining of LC3A, B-II resistant to saponin in (A) CD14^+^ monocytes and (B) lymphocytes. The bars represent the mean values of the percentage of LC3A, B-II cells ± SEM (n = 5). * p <0.05, ** p <0.01. *P* values were calculated using one-way ANOVA and Tukey post hoc multiple comparison test.

It is important to consider that an increase in LC3-II levels can be associated both with a high synthesis of autophagosomes and with a blockage in the autophagic flow (caused by limited fusion of lysosomes or reduced activity of lysosomal enzymes) that reduces the degradation of this protein. To elucidate the origin of LC3-II accumulation versus PGE2 treatment, cells from TB patients were stimulated with *Mtb*-Ag in the presence or absence of the eicosanoid, and incubated with Baf A1 for the last hours of culture. Thus, we were able to corroborate that the addition of Baf A1 causes a significant increase in autophagy levels as compared to monocytes only stimulated with *Mtb*-Ag and treated with PGE2 (Figure 6A). This implies that PGE2 induces the formation of autophagosomes, promoting their maturation and defining a functional autophagic flow. Thus, our findings demonstrate that PGE2 displays an autophagy-inducing effect in human cells. Interestingly, treatment with PGE2 and Baf A1 caused a significant increase in the percentage of LC3A, B-II^+^ lymphocytes as compared to cells only treated with PGE2 (Figure 6B), corroborating an autophagy-inducing effect of PGE2 also in this cell type.

Type I IFN have been described to suppress the production of protective host cytokines after infection with *Mtb*^15^. Furthermore, excessive production of type I IFN is associated with a higher susceptibility to TB^10^. Interestingly, several reports have described that these cytokines display an autophagy-inducing function in different pathological models such as viral infections and cancer^16,17^. However, so far, the contribution of type I interferons to autophagy modulation had not been studied in the context of human TB. Interestingly, IFNα treatment in monocytes stimulated with *Mtb*-Ag did not induce a significant increase in the percentage of CD14^+^ LC3A, B-II^+^ cells (Figure 7A), or in the number of LC3 foci per cell (Figure 7B, C). However, the combination with PGE2 treatment significantly increased autophagy levels measured by flow cytometry and immunofluorescence (Figure 7), reinforcing the promoting role of this lipid mediator in autophagy in human monocytes.

**Figure 7.**
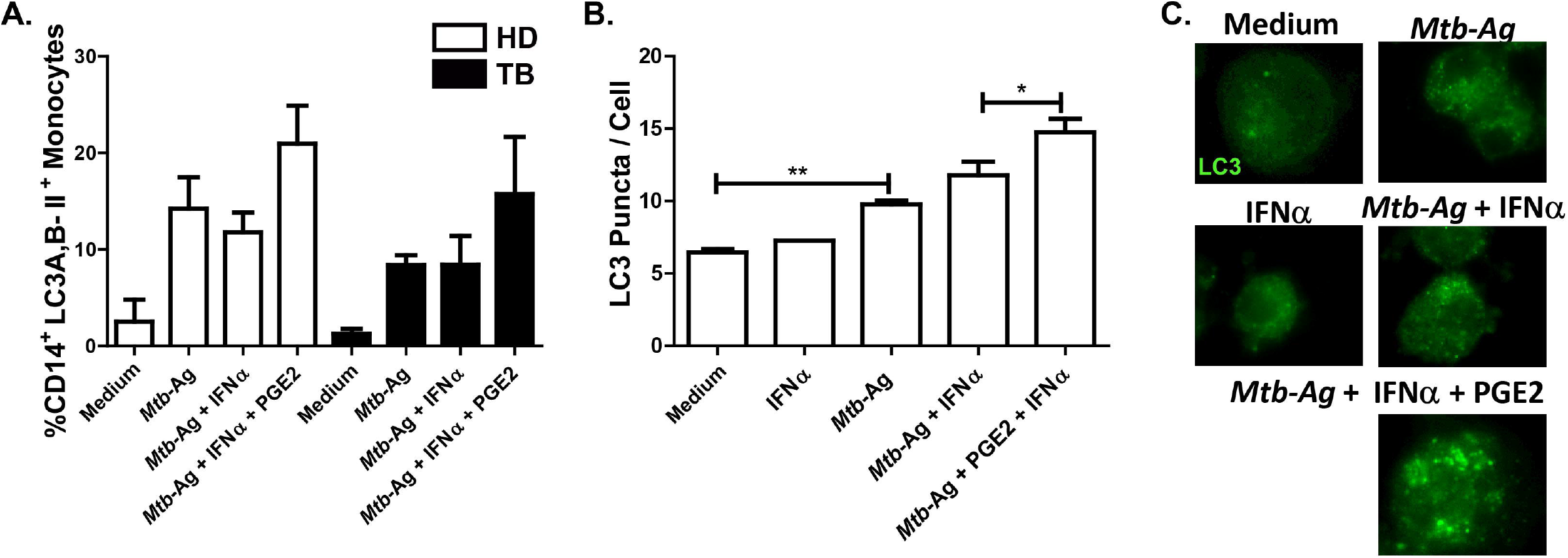
Effect of IFNα on the regulation of autophagy in monocytes from healthy donors and TB patients. **(A-C)** PBMC from HD and TB patients were incubated at 2×10^6^ cells/ml during 16h to allow adherence of monocytes. Afterwards, the cells were stimulated with *Mtb*-Ag (10 μg/ml) in the presence or absence of IFNα (10 ng/ml) or PGE2 (2 μM) during 16h. Autophagy levels were evaluated by **(D)** flow cytometry against intracellular saponin-resistant LC3A,B-II in CD14^+^ cells and by **(E)** immunofluorescence against LC3B in monocytes. **(F)** Representative images of one experiment are shown. Bars represent the mean values of LC3 puncta per cell ± SEM. * p <0.05, ** p <0.01. Statistical differences were calculated using one-way ANOVA and post hoc Dunnett multiple comparison test.

Overall, our findings demonstrate a potent immunosuppressive role for PGE2 during innate and adaptive human immune responses to *Mtb* infection, which could contribute to balance an excessive host response and to the elimination of mycobacteria by promoting autophagic responses.

## Discussion

Even though TB is a curable disease with an efficient and affordable anti-TB treatment, the current prolonged treatment protocols are difficult to maintain, especially in several regions of the world that are severely affected by *Mtb* infection^18^. Host-directed therapies show to provide a largely unexploited approach as adjunctive anti-TB therapies, either directly increasing the ability of the host immune system to eliminate mycobacteria or limiting collateral tissue damage associated with infection. In this regard, it has been suggested that manipulation of PGE2 and/or 5-LO could serve as an approach to counteract the IFN type I response in patients with severe TB^10^, and thus might constitute a host-directed therapy against *Mtb*.

In this work, by analyzing HD and TB patients’ plasma, we found that TB patients exhibit significantly higher concentrations of PGE2 as compared to HD (Figure 1). This observation is in agreement with other works reporting a greater amount of PGE2 in cerebrospinal fluid from meningeal TB patients compared to control individuals^19^. In turn, Mayer-Barber *et al* also observed higher concentrations of PGE2 in plasma from patients with mild TB as compared to HD and, additionally, extended this result taking into account the relationship with the production of another eicosanoid, such as LXA-410.

Given the multiple functions of PGE2 in various cell types, and throughout different stages of the immune response, this eicosanoid presents the paradox of being a proinflammatory factor with immunosuppressive activity. In terms of the latter, we observed that PGE2 treatment inhibited the proliferation of *Mtb*-stimulated T cells from HD and TB patients (Figure 2). In fact, treatment with indomethacin, a selective COX-2 inhibitor, increases the lymphocyte proliferation in HD and TB and leprosy patients^20^. Strikingly, Tonby *et al* reported that the use of this inhibitor decreases the proliferation and cytokine secretion of mycobacterial antigen stimulated lymphocytes from TB patients. However, these observations could be linked to a direct inhibitory effect of indomethacin on the activation of the NF-κB pathway^21^.

Optimal T cell activation requires costimulatory signaling provided by the interaction of molecules expressed on the antigen presenting cell with specific ligands on the T lymphocyte. Therefore, we also study the effect of PGE2 on the expression of different on T lymphocytes and monocytes. We observed that 5 days treatment with PGE2 significantly decreased the expression of SLAMF1 and CD31 on CD4^+^ T lymphocytes from HD and TB patients. Accordingly, we previously demonstrated that costimulation through SLAMF1 increases IFNγ expression in cells from TB patients^22,23^. In turn, CD31, also known as PECAM-1, has been reported to facilitate transendothelial leukocyte migration^24^. However, our laboratory has shown that the interaction between CD31 and SAP proteins negatively regulates the pathways that lead to IFNγ production during active pulmonary TB^25^. Additionally, a reduction in the expression of the costimulatory molecule CD80 and MHC class I molecules was found on CD14^+^ monocytes treated with PGE2 (Figure 3). These results demonstrate that PGE2 during stimulation with *Mtb*-Ag downregulates the expression of various membrane receptors involved in T cell activation. In this regard, the addition of PGE2 has been shown to reduce the expression of another costimulatory molecule, CD46 on activated T cells^26^. Furthermore, PGE2 also inhibits the expression of CD80, CD86 and MHC class I after LPS stimulation^27^. However, in other models, it was reported that PGE2 might increase dendritic cell maturation^28^, denoting the importance of the context, exposure time and concentration of this eicosanoid.

Moreover, we observed that PGE2 treatment significantly reduced IFNγ and TNFα secretion from *Mtb*-Ag stimulated PBMC (Figure 4). Thus, our data indicate that PGE2 affected the production of critical cytokines required in the immune response against this pathogen. Overall, these results suggest a clear immunosuppressive effect of elevated PGE2 during the human immune response against *Mtb*.

However, it remains to be understood how this lipid mediator can collaborate with the host response to contain the pathogen as suggested before. Protection against necrosis (by preventing damage to the internal mitochondrial membrane^5^ and promoting rapid repair of the plasma membrane^7^) and the switch towards programmed cell death by apoptosis are key mechanisms in the resolution of the infection. We and other authors have demonstrated that activation of autophagy can result in the elimination of the bacteria in autolysosomes^14,29,30^. Furthermore, autophagy also controls inflammation by regulating signaling pathways of the innate immunity, removing endogenous agonists from the inflammasome and by modulating the secretion of immune mediators. For these reasons, we next evaluated whether PGE2 could be contributing to an autophagic response during infection with *Mtb*.

Interestingly, using flow cytometric and confocal microscopy approaches, we demonstrated that treatment with PGE2 induced autophagy in PBMC stimulated with *Mtb*-Ag. Furthermore, the exogenous addition of this eicosanoid induced a functional autophagy flow both in monocytes and lymphocytes from HD and TB patients (Figure xx). In different pathological models, including tenofibroblasts and human colon cancer cells, the role of PGE2 in modulating autophagy has been demonstrated^11,12,31^. Furthermore, inhibition of COX-2 has been shown to reverse autophagy induced during lupus nephritis^32^. Besides, kidney autophagy levels were shown to decrease after silencing PGES-2^33^. Importantly, Xiong *et al* reported that COX-2 suppresses mycobacterial growth in murine macrophages by promoting autophagy via the AKT/mTOR pathway^13^. Strikingly, other authors described that the PGE2 reverses vitamin D3-induced autophagy in macrophages infected with *Mtb*^34^. Nevertheless, this work requires a better evaluation of autophagy flow in the studied cells.

Interestingly, we detected increased autophagy in lymphocytes stimulated with mycobacterial antigens in the presence of PGE2 (Figure 5B). Numerous investigations have shown that autophagy is essential for CD4^+^ T cell survival and homeostasis in peripheral lymphoid organs. Moreover, autophagy is also required for T cell proliferation and cytokine production in response to TCR activation^35^. Besides, autophagy is activated in regulatory T cells and supports lineage stability and survival^36^. Accordingly, we previously demonstrated that PGE2 contributes to the expansion of regulatory T cells in response to mycobacterial antigens^9^. Thus, PGE2 activation of autophagy in lymphocytes might directly participate in the maintenance of these cells, and could indirectly act in the generalized immunosuppression observed.

In different models of viral infections, type I IFNs have been found to confer protection against these intracellular pathogens by inducing autophagy^37^. However, most studies support the findings that type I IFNs actually promote infection by *Mtb*^38,39^. Antonelli *et al* have shown that IFN receptor type I deficient mice chronically infected with several *Mtb* strains showed reduced bacterial loads compared to WT infected animals^40^. Our analyses by flow cytometry and confocal microscopy revealed that TB patients’ monocytes stimulated with *Mtb*-Ag in the presence of IFNα did not modify autophagy levels (Figure 7). However, exogenous addition of PGE2, even under these conditions of high concentrations of type I IFN, significantly induced autophagy in those cells (Figure 7). This finding reinforces the inductor role of this lipid mediator in monocyte autophagy, especially considering that type I IFNs have been described to promote an IFN-refractory macrophage phenotype that inhibits autophagy induced by the latter cytokine and stimulates mycobacteria growth^41^.

In summary, our findings suggest that PGE2 would have a main suppressive role during innate and adaptive immunity against *Mtb*, participating in homeostasis of this response and protecting the host from excessive inflammation. Moreover, we hypothesized that PGE2-induced autophagy could be involved in immunity suppression and bacterial clearance. Rangel Moreno *et al* have described a differential participation of this lipid mediator during the early and late phases of experimental pulmonary TB^8^. In conclusion, the outcome of PGE2 inhibition or stimulation might vary during the acute and chronic stages of tuberculosis infection. The design of new personalized treatments focused on the regulation of host’s eicosanoids will have to consider these various consequences.

## Material and Methods

### Subjects

HIV-uninfected patients with TB were diagnosed at the Servicio de Tisioneumonología Hospital F.J. Muñiz, Buenos Aires, Argentina, based on clinical and radiological data, together with the identification of acid-fast bacilli in sputum. All participating patients had received less than 1 week of anti-TB regular therapy. Bacillus Calmette-Guerin (BCG)-vaccinated healthy control individuals (HD) from the community participated in this study. Peripheral blood was collected in heparinized tubes from each participant after obtaining informed consent. The local Ethics Committee approved all studies.

### Exclusion Criteria

All subjects were 18-60 years old and had no history of illnesses that affect the immune system, such as HIV infection, a recent diagnosis of cancer, treatment with immunosuppressive drugs, hepatic or renal disease, pregnancy, or positive serology for other viral (e.g., hepatitis A, B or C), or bacterial infections (e.g., leprosy, syphilis). Individuals with bleeding disorders or under anticoagulant medication that might be at an increased risk of bleeding during the procedure of obtaining the sample were excluded from the study. Individuals with latent infection were excluded of the present study by using the QuantiFERON-TB^®^ GOLD PLUS (Qiagen, 622120 and 622526).

### Plasmatic PGE2 determination

Plasma samples were collected from heparinized peripheral blood from HD and TB patients, centrifuged for 15 minutes at 1000 x g at 4 ° C. They were stored at −70°C. PGE2 measurement was performed by radioimmunoassay (RIA) as previously^42^. Radioactivity was measured in a beta scintillation counter. After a logarithmic transformation the data was expressed as pg of PGE2/ml of plasma. The method has a cross reactivity of less than 0.1% and a sensitivity of 5 pg/tube with a Ka = 1.5 x 10^10^/ mol.

### Antigen

*In vitro* stimulation of cells was performed with a cell lysate from the virulent *Mycobacterium tuberculosis* strain H37Rv, prepared by probe sonication (*Mtb*-Ag) (BEI Resources, NIAID, NIH: *Mycobacterium tuberculosis*, Strain H37Rv, whole cell lysate, NR-14822).

### Cell preparations and culture conditions

Ficoll-Hypaque (GE Healthcare, 17-1440-03) and cultured (2×10^6^ cells/mL) in flatbottom 24 or 48-well plates with RPMI 1640 (Invitrogen, 22400071) supplemented with L-Glutamine (2mM, Sigma), Penicillin/Streptomycin, and 10% Fetal Bovine Serum (FBS; Gibco, 10437028) for 16 h without stimulus to allow monocyte adherence. Cells were then stimulated with sonicated *Mtb* (*Mtb*-Ag, BEI Resources, NIH, 10 μg/ml) ± PGE2 (2 uM, Sigma, P0409) ± IFNα 2a (10 ng/ml, Biosidus) for different time points. In order to determine the effect of different treatments on the autophagic flux, the vacuolar-type HC-ATPase inhibitor bafilomycin A1 (100 nM; Fermentek, 88899-55-2) was added for the last 2 h of culture before LC3 determination by flow cytometry.

### Proliferation assay

PBMC were stimulated with *Mtb*-Ag for five days in the presence or absence of PGE2. Cells were pulsed with [^3^H]TdR (1 μCi/well, Perkin Elmer, MA, EE.UU) and harvested 16h later. [^3^H]TdR incorporation (c.p.m.) was measured in a liquid scintillation counter (Wallac 1214 Rackbeta, Turku, WF, Finland. Proliferation index for each individual was calculated as cpm after *Mtb*-Ag stimulation/cpm after culturing with medium

### Flow Cytometry

To determine the level of immune receptors expression on lymphocytes and monocytes, PBMC stimulated with *Mtb*-Ag (10 μg/ml) treated or not with PGE2 (2 mM) were blocked in PBS (137 mM NaCl, 2.7 mM KCl, 8 mM Na_2_HPO_4_, and 2 mM KH_2_PO_4_)-FBS 5% for 15 min and then stained for surface expression with fluorophore-marked antibodies against SLAMF1 (BD, 559572), CD31 (BD, 560984), CD80 (Biolegend, 305207), MHC-I (Biolegend, HLA-A2, 343306), MHC-II (Biolegend, HLA-DQ, 318106), CD3 (Biolegend, 300306) and CD14 (Biolegend, 367116).

Intracellular staining of endogenous saponin-resistant LC3 was performed as described^14^. Briefly, cells were washed with PBS and then permeabilized with PBS containing 0.05% saponin. In this protocol, the cells are not fixed, therefore LC3-I is washed out of the cell because, unlike LC3-II, it is not anchored to the autophagosome. Cells were then incubated with mouse anti-human LC3A/B antibody (MBL International, M152-3) for 20 min, rinsed with PBS, incubated with anti-mouse secondary antibody conjugated to fluorescein isothiocyanate (eBioscience,62-6511) for 20 min and rinsed twice with PBS. Afterwards, cells were stained with anti-CD14 or anti-CD3 antibodies (Biolegend, 325608 and 300308) to detect the monocyte and T lymphocytes populations.

Negative control samples were incubated with irrelevant isotype matched monoclonal antibody (Biolegend, 400140). Samples were analyzed on a FACSAria II flow cytometer (BD Biosciences).

### ELISA

Culture supernatants of PBMC stimulated or not with *Mtb*-Ag in the presence or absence of PGE2 were obtained to evaluate cytokine levels by ELISA. TNFα, IL-1β and IFNγ (BioLegend) secretion was measured in by ELISA following the manufacturers’ instructions.

### Confocal microscopy

Cells were cultured and stimulated or infected on coverslips for 16 h. After incubation under different experimental conditions, cells were washed in order to remove nonadherent cells. Adherent cells were then fixed with cold methanol for 20 sec, then washed and subsequently permeabilized and blocked with blocking buffer (PBS containing 0.5% saponin (Santa Cruz Biotechnology, sc-280079A) and 1% bovine serum albumin (Santa Cruz Biotechnology, sc-2323A) for 15 min. The buffer was then removed and the LC3 primary antibody was added (Cell Signaling Technology, 2775) and incubated for 16 h at 4° C. Afterwards, cells were washed with blocking buffer and incubated with the secondary antibody (Alexa Fluor^®^ 488 Goat Anti-Rabbit IgG (HCL); Invitrogen, A11008) for 2 h at room temperature. Finally, nuclei were stained with DAPI. The coverslips were mounted with PBS-glycerol (Sigma-Aldrich, G2025) and fixed cells were imaged using a Zeiss Spectral LSM 510 confocal microscope (Zeiss, Jena, Germany) using objective 63, numerical aperture (NA) 1.42.

### Image processing

All the images were processed using ImageJ software (Wayne Rasband, National Institutes of Health). After the image binarization using a defined threshold, the number of LC3 puncta was quantified using the Particle Analyzer plugin. Brightness and contrast were adjusted in all images belonging to the same individual, when needed.

### Statistical Analysis

Analysis of variance and post hoc Tukey’s multiple comparisons test were used as indicated in the figure legend. Mann–Whitney *U* test and Wilcoxon rank sum test were used for the analysis of unpaired and paired samples respectively. *P* values of < 0.05 were considered statistically significant.

## Acknowledgments

We are thankful to Licenciado Guillermo Piazza and Dr. Julieta Schander for their constant support and technical assistance.

## References

1. Daniel, T. M. The history of tuberculosis. Respir. Med. 100, 1862–1870 (2006).

2. Discovery and preclinical development of new antibiotics. https://www.tandfonline.com/doi/full/10.3109/03009734.2014.896437.

3. WHO | Global tuberculosis report 2019. https://www.who.int/tb/publications/global_report/en/.

4. Dietzold, J., Gopalakrishnan, A. & Salgame, P. Duality of lipid mediators in host response against Mycobacterium tuberculosis: good cop, bad cop. F1000prime Rep. 7, 29 (2015).

5. Chen, M. et al. Lipid mediators in innate immunity against tuberculosis: opposing roles of PGE2 and LXA4 in the induction of macrophage death. J. Exp. Med. 205, 2791–2801 (2008).

6. An important role of prostanoid receptor EP2 in host resistance to Mycobacterium tuberculosis infection in mice - PubMed. https://pubmed.ncbi.nlm.nih.gov/23033144/.

7. Divangahi, M. et al. Mycobacterium tuberculosis evades macrophage defenses by inhibiting plasma membrane repair. Nat. Immunol. 10, 899–906 (2009).

8. The role of prostaglandin E2 in the immunopathogenesis of experimental pulmonary tuberculosis. https://www.ncbi.nlm.nih.gov/pmc/articles/PMC1782721/.

9. Garg, A. et al. MANNOSE-CAPPED LIPOARABINOMANNAN- AND PROSTAGLANDIN E2-DEPENDENT EXPANSION OF REGULATORY T CELLS IN HUMAN Mycobacterium tuberculosis INFECTION. Eur. J. Immunol. 38, 459–469 (2008).

10. Mayer-Barber, K. D. et al. Host-directed therapy of tuberculosis based on interleukin-1 and type I interferon crosstalk. Nature 511, 99–103 (2014).

11. Chen, H., Chen, L., Cheng, B. & Jiang, C. Cyclic mechanical stretching induces autophagic cell death in tenofibroblasts through activation of prostaglandin E2 production. Cell. Physiol. Biochem. Int. J. Exp. Cell. Physiol. Biochem. Pharmacol. 36, 24–33 (2015).

12. Pelissier-Rota, M. A., Pelosi, L., Meresse, P. & Jacquier-Sarlin, M. R. Nicotine-induced cellular stresses and autophagy in human cancer colon cells: A supportive effect on cell homeostasis via up-regulation of Cox-2 and PGE2 production. Int. J. Biochem. Cell Biol. 65, 239–256 (2015).

13. Xiong, W. et al. Novel Function of Cyclooxygenase-2: Suppressing Mycobacteria by Promoting Autophagy via the Protein Kinase B/Mammalian Target of Rapamycin Pathway. J. Infect. Dis. 217, 1267–1279 (2018).

14. Tateosian, N. L. et al. IL17A augments autophagy in Mycobacterium tuberculosis-infected monocytes from patients with active tuberculosis in association with the severity of the disease. Autophagy 13, 1191–1204 (2017).

15. McNab, F., Mayer-Barber, K., Sher, A., Wack, A. & O’Garra, A. Type I interferons in infectious disease. Nat. Rev. Immunol. 15, 87–103 (2015).

16. Schmeisser, H., Bekisz, J. & Zoon, K. C. New function of type I IFN: induction of autophagy. J. Interferon Cytokine Res. Off. J. Int. Soc. Interferon Cytokine Res. 34, 71–78 (2014).

17. Zhao, J. et al. Interferon-alpha-2b induces autophagy in hepatocellular carcinoma cells through Beclin1 pathway. Cancer Biol. Med. 11, 64–68 (2014).

18. The Lancet Respiratory Medicine Commission: 2019 update: epidemiology, pathogenesis, transmission, diagnosis, and management of multidrug-resistant and incurable tuberculosis - The Lancet Respiratory Medicine. https://www.thelancet.com/journals/lanres/article/PIIS2213-2600(19)30263-2/fulltext.

19. The prevalence, characteristics and outcome of seizure in tuberculous meningitis | SpringerLink. https://link.springer.com/article/10.1186/s42494-020-0010-x.

20. Bahr, G. M., Rook, G. A. & Stanford, J. L. Prostaglandin-dependent regulation of the in vitro proliferative response to mycobacterial antigens of peripheral blood lymphocytes from normal donors and from patients with tuberculosis or leprosy. Clin. Exp. Immunol. 45, 646–653 (1981).

21. Tonby, K. et al. The COX-inhibitor indomethacin reduces Th1 effector and T regulatory cells in vitro in Mycobacterium tuberculosis infection. BMC Infect. Dis. 16, 599 (2016).

22. Pasquinelli, V. et al. IFN-γ Production during Active Tuberculosis Is Regulated by Mechanisms That Involve IL-17, SLAM, and CREB. J. Infect. Dis. 199, 661–665 (2009).

23. Quiroga, M. F. et al. Activation of signaling lymphocytic activation molecule triggers a signaling cascade that enhances Th1 responses in human intracellular infection. J. Immunol. Baltim. Md 1950 173, 4120–4129 (2004).

24. Muller, W. A. Leukocyte-endothelial-cell interactions in leukocyte transmigration and the inflammatory response. Trends Immunol. 24, 327–334 (2003).

25. Cross-Talk between CD31 and the Signaling Lymphocytic Activation Molecule— Associated Protein during Interferon-γ Production against Mycobacterium tuberculosis | The Journal of Infectious Diseases | Oxford Academic. https://academic.oup.com/jid/article/196/9/1369/2191997.

26. Kickler, K. et al. Prostaglandin E2 affects T cell responses through modulation of CD46 expression. J. Immunol. Baltim. Md 1950 188, 5303–5310 (2012).

27. Jung, I. D. et al. COX-2 and PGE2 signaling is essential for the regulation of IDO expression by curcumin in murine bone marrow-derived dendritic cells. Int. Immunopharmacol. 10, 760–768 (2010).

28. Regulation of Immune Responses by Prostaglandin E2. https://www.ncbi.nlm.nih.gov/pmc/articles/PMC3249979/.

29. Castillo, E. F. et al. Autophagy protects against active tuberculosis by suppressing bacterial burden and inflammation. Proc. Natl. Acad. Sci. 109, E3168–E3176 (2012).

30. Autophagy Is a Defense Mechanism Inhibiting BCG and Mycobacterium tuberculosis Survival in Infected Macrophages: Cell. https://www.cell.com/fulltext/S0092-8674(04)01106-7.

31. Prostaglandin E2 as a Regulator of Immunity to Pathogens. https://www.ncbi.nlm.nih.gov/pmc/articles/PMC5898978/.

32. Jin, J. et al. Activation of Cyclooxygenase-2 by ATF4 During Endoplasmic Reticulum Stress Regulates Kidney Podocyte Autophagy Induced by Lupus Nephritis. Cell. Physiol. Biochem. Int. J. Exp. Cell. Physiol. Biochem. Pharmacol. 48, 753–764 (2018).

33. Aggravation of acute kidney injury by mPGES-2 down regulation is associated with autophagy inhibition and enhanced apoptosis | Scientific Reports. https://www.nature.com/articles/s41598-017-10271-8.

34. Wan, M. et al. Prostaglandin E2 suppresses hCAP18/LL-37 expression in human macrophages via EP2/EP4: implications for treatment of Mycobacterium tuberculosis infection. FASEB J. Off. Publ. Fed. Am. Soc. Exp. Biol. 32, 2827–2840 (2018).

35. Cell-Intrinsic Roles for Autophagy in Modulating CD4 T Cell Functions. https://www.ncbi.nlm.nih.gov/pmc/articles/PMC5954027/.

36. Autophagy in regulatory T cells: A double-edged sword in disease settings - ScienceDirect. https://www.sciencedirect.com/science/article/pii/S0161589018308575.

37. Crosstalk between Autophagy and Type I Interferon Responses in Innate Antiviral Immunity. https://www.ncbi.nlm.nih.gov/pmc/articles/PMC6409909/.

38. Flynn, J. L. & Chan, J. What’s good for the host is good for the bug. Trends Microbiol. 13, 98–102 (2005).

39. TPL-2–ERK1/2 Signaling Promotes Host Resistance against Intracellular Bacterial Infection by Negative Regulation of Type | IFN Production | The Journal of Immunology. https://www.jimmunol.org/content/191/4/1732.

40. Antonelli, L. R. V. et al. Intranasal Poly-IC treatment exacerbates tuberculosis in mice through the pulmonary recruitment of a pathogen-permissive monocyte/macrophage population. J. Clin. Invest. 120, 1674–1682 (2010).

41. Lienard, J., Movert, E., Valfridsson, C., Sturegård, E. & Carlsson, F. ESX-1 exploits type | IFN-signalling to promote a regulatory macrophage phenotype refractory to IFNγ-mediated autophagy and growth restriction of intracellular mycobacteria. Cell. Microbiol. 18, 1471–1485 (2016).

42. Ribeiro, M. L. et al. Epidermal growth factor prevents prepartum luteolysis in the rat. Proc. Natl. Acad. Sci. 102, 8048–8053 (2005).

